# Metagenome-based vertical profiling of the Gulf of Mexico highlights its uniqueness and far-reaching effects of freshwater input

**DOI:** 10.1101/2025.07.17.665386

**Authors:** Roth E. Conrad, Despina Tsementzi, Alexandra Meziti, Janet K. Hatt, Joseph Montoya, Konstantinos T. Konstantinidis

## Abstract

Genomic and metagenomic explorations of the oceans have identified well-structured microbial assemblages showing endemic genomic adaptations with increasing depth. However, deep water column surveys have been limited, especially of the Gulf of Mexico (GoM) basin, despite its importance for human activities. To fill this gap, we report on 19 deeply sequenced [∼5 Gbp/sample] shotgun metagenomes collected along a vertical gradient, from the surface to about 2,000m deep, at three GoM stations. Beta diversity analysis revealed strong clustering by depth, and not by station, even when including previously determined samples from other ocean basins. However, a community level pangenome style gene content analysis revealed ∼54% of predicted gene sequences to be station specific within our GoM samples. Of the 154 high-quality MAGs recovered, 145 represent novel species compared to NCBI genomes and TARA MAGs databases. Two of these MAGs were relatively abundant at both surface and deep samples, revealing remarkable versatility across the water column. A few MAGs of freshwater origin (∼6% of total detected) were relatively abundant at 600m deep and 270 miles from the coast at one station, revealing that the effects of freshwater input in the GoM can sometimes be far-reaching and long-lasting. Notably, 1,447/16,068 of the total COGs detected were positively (Pearson’s r ≥ 0.5) or negatively (Pearson’s r ≤ −0.5) correlated with depth including beta-lactamases, dehydrogenase, and CoA-associated oxidoreductases. Taken together our results reveal substantial novel genome and gene diversity across the GoM’s water column, and testable hypotheses for some of the diversity patterns observed.

**IMPORTANCE:** To what extent microbial communities are similar between different ocean basins at similar depths and what the impact of freshwater input by major rivers may be on these communities remain poorly understood issues with potentially important implications for modeling and managing marine biodiversity. In this study, we performed metagenomic sequencing and recovered 154 high-quality metagenome-assembled genomes (MAGs) from three stations in the Gulf of Mexico (GoM) and from various depths up to about 2000m. Comparison to similar data from other ocean basins highlight the unique diversity harbored by the GoM, which could be driven by more substantial input by the Mississippi River and by human activities, including offshore oil drilling. The data and results provided by this study should be useful for future comparative analysis of marine biodiversity and contribute toward its more complete characterization.

## INTRODUCTION

While an increasing part of extant microbial diversity is being discovered via (mostly) culture-independent metagenomics approaches, the majority of the diversity has not been recovered yet, especially for enormous environments such as the oceans (1–5). Cataloguing this diversity is important for several reasons, including to better understand how this powerful biogeochemical force may be affected in dynamic and changing world (6–11). To this end, efforts have been undertaken in the past two decades to collect metagenomic samples that cover representative portions of the global ocean (12–19). However, since additional insights are still being gained from every new sample, more sampling is needed across space and time, particularly in regions such as the Gulf of Mexico (GoM) that are important to and heavily influenced by human activities yet remain under sampled.

Since the first microbial sequences from the surface and deep ocean were compared, genetic differences have been reported between the two habitats (20, 21). For instance, several studies have revealed differences in genome size and amino acid composition between the photic zone and the abyss (22). With increased genome size with increasing depth, the abundance of certain gene functions like transposases and integrases have also been noted to increase with depth (23–25). The physicochemical differences between the surface and deep ocean including light intensity, salinity, temperature, pressure, and nutrient availability presumably drive some of these genetic adaptations. However, to better quantify and understand the effects of these physicochemical parameters and their distribution throughout different ocean basins, more vertical water column profiles are clearly needed, especially below the photic zone.

Specifically, few metagenomes have been sequenced from the GoM and these were typically associated with oil spills, hydrocarbon seeps and hypoxic zones (26–31). In this study, samples were collected during a research voyage targeting surface *Trichodesmium* blooms, which were identified and tracked via satellite imaging, during a period without major oil inputs. Accordingly, our study enabled the sampling of natural bloom events across spatial gradients. Additionally, we aim to assess how the unique characteristics and geography of the GoM— including substantial riverine inputs from the Mississippi River and others—influence the vertical stratification of microbial assemblages.

To provide additional metagenomic perspective on the ocean water column, we sequenced 15 shotgun metagenomes in the northern GoM at five distinct depths ranging from the surface down to the oxygen minimum zones (OMZ) between 300-500m. We sequenced 4 additional shotgun metagenomes at one station reaching below the OMZ down to 2,100m. We analyzed microbial community diversity and taxonomic composition as well as genetic similarity and functional gene abundance in each sample. We report the fraction of common gene sequences shared between samples compared at 40% and 70% amino acid sequence identity (roughly corresponding to family and genus levels, respectively) as well as 90% and 95% nucleotide sequence identity (species level). We used two thresholds for the species level in order to account for the most frequently observed area of discontinuity among species at 95% genome-aggregate average nucleotide identity (ANI) (32) (95% threshold used), but also the facts that several abundant marine taxa like SAR11 show high intraspecies diversity [area of discontinuity is at 90-91% ANI (33)] and fast-evolving genes in the genome may sometimes show lower identity values than 95% (90% threshold used). We also report gene copy counts and relative abundance of predicted functions in each sample and describe the functional gene content that correlates well with ocean density (which correlates with temperature and depth). Finally, we present novel GoM associated metagenome assembled genomes (MAGs) analyzed with the Microbial Genome Atlas (MiGA) alongside their relative abundance distribution across our sample set.

## RESULTS

### Study Stations

Water column samples were collected during CTD casts from the May 2012 R/V Endeavour cruise EN509 and shotgun metagenomes were sequenced from the surface (∼3m), the mixed layer (ML; 15-25m), the deep chlorophyll maximum (DCM; 70-90m), below the DCM but above the oxygen minimum zone (aOMZ; 100-150m), and the OMZ (200-400m) from three stations (2, 5, and 8) in the northwest GoM (Fig. 1A). Four additional metagenomes were sequenced from below the OMZ (Deep; 600, 1000, 1470 and 2107m) from station 5. Station 2 was near the edge of the Texas-Louisiana Shelf about 50 miles southeast from the mouth of the Mississippi river. Station 8 and 5 were over the TX-LA Slope about 190 and 270 mi. southeast of Galveston, TX with station 8 nearer the TX-LA Shelf and station 5 nearer the edge of the TX-LA slope. Fluorescence, oxygen, salinity, and temperature measurements were similar at all stations except salinity was lower (29.6 vs. 36 PSU) and fluorescence was increased (0.9 vs. 0.07 mg/m^3^) at the surface of station 2 while the fluorescence peak of the DCM layer was reduced (0.3 vs. 0.8 mg/m^3^) compared to stations 5 and 8 (Fig. 1B, C, D, and E). Temperature and salinity were most similar for all samples from the ML and DCM while salinity was variable across surface samples and temperature and salinity were variable across aOMZ and OMZ samples (Fig. 1E).

**Figure 1.**
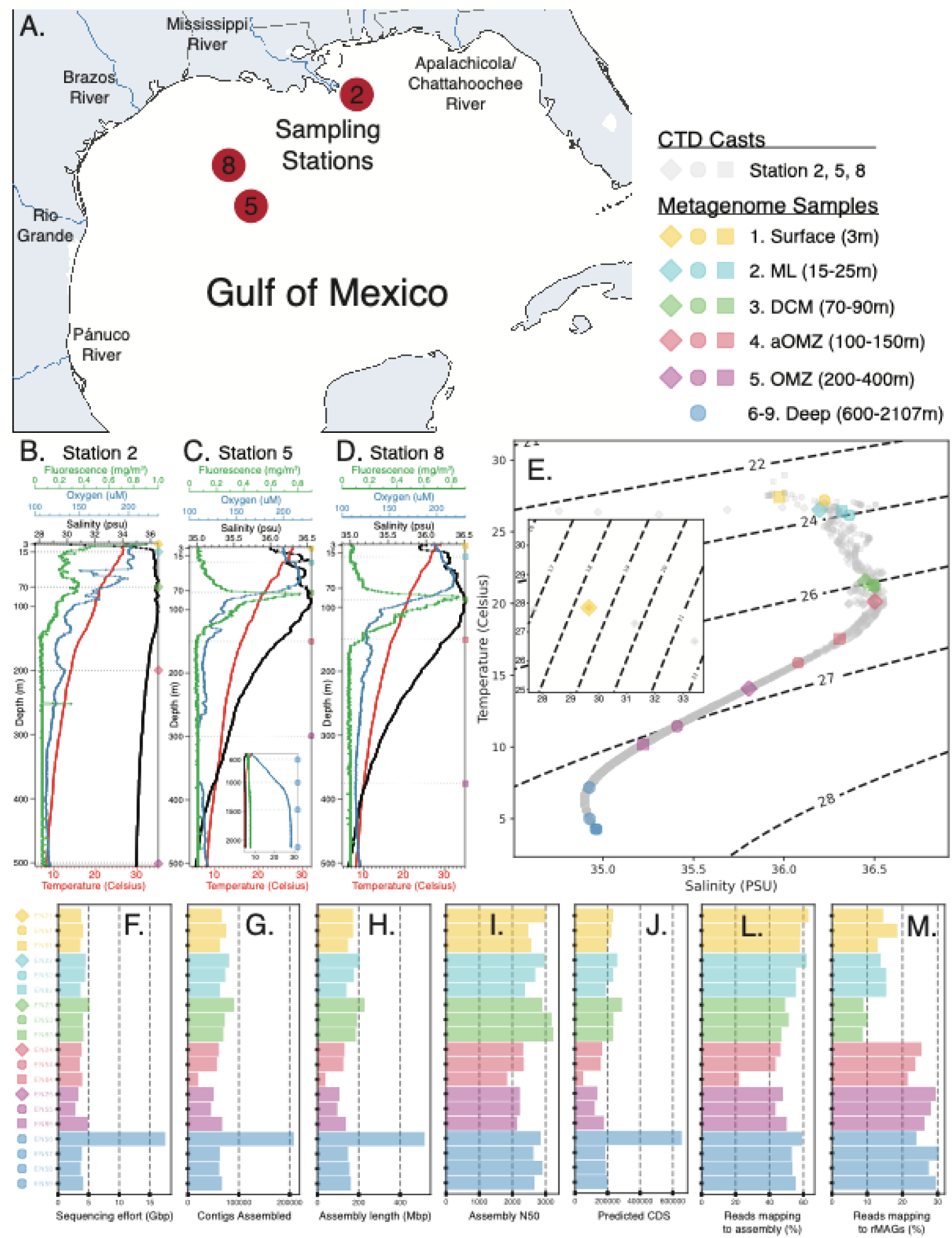
Overview of samples sequenced for this study. CTD casts were made at three stations in the Gulf of Mexico (panel A) and temperature, salinity, oxygen, and fluorescence measurements were recorded along the depth gradient of the water column for Station 2 (B), Station 5 (C), and Station 8 (D). Temperature-salinity diagram of metagenome sample collection points with isopycnic lines expressing density as **σt** = (ρ-1000)[kg/m^3] (E). Stations are marked with diamonds (Station 2), circles (Station 5), or squares (Station 8). Colors denote the depth layer (1–9) of metagenome sample collection. Sequencing effort (F), the number of contigs assembled (G), total assembly length in megabasepairs (H), assembly N50 (I), predicted CDS (J), unassembled reads mapping back to the assembled contigs (L), and unassembled reads mapping to rMAGs (M) are reported for each metagenome sample analyzed.

### Metagenome Statistics

Sequencing effort, except for sample EN56 which was sequenced about 4 times more deeply, ranged from 3-5 Gbp (giga base pairs) for all samples (Fig. 1F), and covered an average of 58% of the estimated sequence diversity of the microbial communities sampled (Fig. 2C; Sup. File 1). The diversity covered by sequencing effort decreased with depth (Pearson’s r = −0.32; Fig. 2C; Sup. File 1, GoM DivCov tab), due presumably to increased genome sizes and greater microbial diversity observed in deeper samples (see also below). The number of contigs assembled, the overall assembly length in Mbp (mega base pairs), and the predicted coding sequences (CDS) were moderately correlated with sequencing effort (Pearson’s r ∼ 0.6) while assembly N50 was only weakly correlated (Pearson’s r = 0.31; Fig. 1F-J; Sup. File 1, GoM Effort tab). Excluding sample EN56, an average of approximately 64,500 contigs and 195,000 CDS were recovered per sample. The 340% increase in sequencing effort for sample EN56 yielded over 144,000 more contigs and 460,000 more CDS on average compared to the other samples (224-237% increase; Fig. 1F-J). Nonetheless, reads used in the assembly and the diversity covered by sequencing effort remained within the 22-63% (Fig. 1L) and 47-77% (Fig. 2C) range of the other samples, respectively, which showed that while greater sequencing effort increased DNA and gene recovery, the amount of recovered but unassembled DNA sequence and rare biosphere sequences also increased (Sup. File 1). Only 22-50% of the sequenced reads were used by the assembled contigs across samples, and only 9-30% were mapped back to the representative metagenome assembled genomes (rMAGs) binned from those contigs (Fig 1L-M; Sup. File 1), revealing that an average of 52% of the total diversity captured by sequencing was represented by the assembled sequences and predicted CDS.

**Figure 2.**
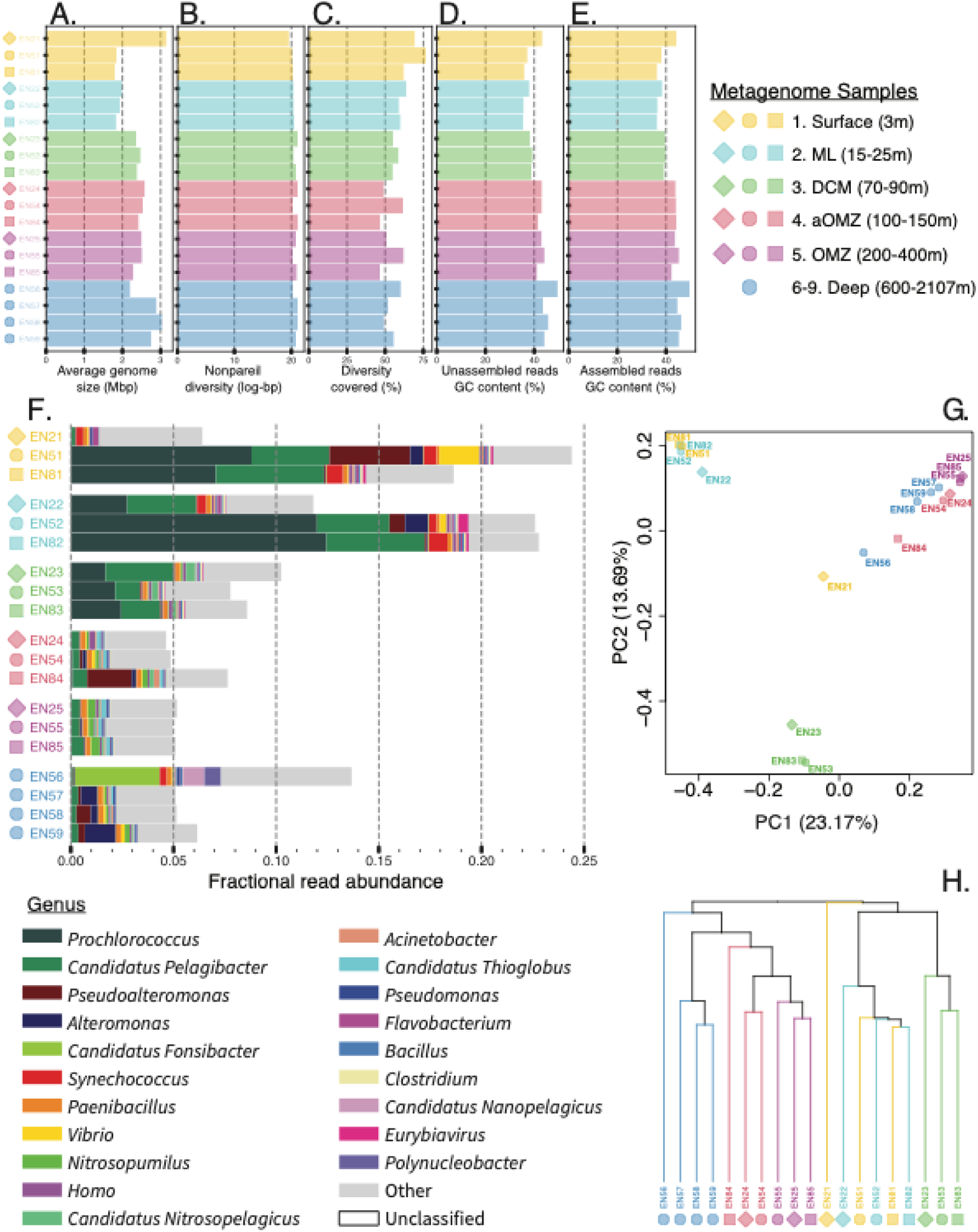
Community level diversity metrics. Average genome size estimates from MicrobeCensus (A), Alpha diversity estimates from Nonpareil (B), Diversity covered with sequencing effort estimates by Nonpareil (C), and GC content of unassembled (D) vs assembled (E) reads are reported for each sample. The taxonomic distribution of the top 20 most abundant genus in each sample as reported by Kraken with Bracken (E). Note the overwhelming fraction of unclassified reads (reads with no match to the current databases). Beta diversity as assessed by Simka jaccard distance shown as a PCoA (G) or hierarchical clustering (H). Note strong sample grouping by depth layer.

While the reads represented by assembled contigs from the surface and ML samples were in the upper range of all samples (56-63%), the reads represented by rMAGs from samples below the DCM were significantly greater (p < 0.05) than in the DCM and above samples (Fig. 1. L-M; Sup. File 1, GoM Effort tab), which suggested that the deeper water communities are better covered by rMAGs in our analysis. Correspondingly, there were 73 more rMAGs recovered from the aOMZ and below (68 rMAGs surface – DCM vs. 141 rMAGs aOMZ – Deep; Sup. File 4, MAG-Sample tab) which seemed to explain this discrepancy in rMAG read representation above and below the DCM and indicated better MAG recovery from samples below the DCM. However, 65 of the 141 rMAGs recovered from below the DCM originated from sample EN56, and thus, better MAG yield was also associated with the increased sequencing depth from this sample. Even so, after excluding sample EN56, there is still an 8 MAG difference below the DCM and the mean difference of rMAG recovery per sample is still significant (76 rMAGs below DCM – 68 rMAGs above; p < 0.05; Sup. File 4, MAG-Sample tab) which yielded a 13.4% increase in community representation by rMAGs based on read mapping results below the DCM (Sup. File 1, GOM Effort tab). Regardless, the G+C% content of the assembled contigs remained similar (mean difference < 1%) to that of the unassembled reads suggesting that the assembly is likely not biased toward any specific proportion of the community and is likely representative, at least in terms of G+C% (Fig. 2D-E; Sup. File 1).

### Microbial Community Metrics

Apart from sample EN21 which was an outlier, average genome size had the strongest correlation with density (Pearson’s r = 0.86) followed by temperature (Pearson’s r = −0.80) and depth (Pearson’s r = 0.67), which indicated microbial genome size increased in denser, colder, and deeper water masses (Fig. 2A; Sup. File 1, GoM GenomeSize tab). Likewise, G+C% content also increased (excluding EN21) most strongly with density (Pearson’s r = 0.88) followed by temperature (Pearson’s r = −0.87) and depth (Pearson’s r = 0.58; Sup. File 1, GoM %GC tab). While alpha diversity was only weakly correlated with depth across all samples (Pearson’s r = 0.37), alpha diversity correlation with depth was stronger when each sampling station was considered separately (Pearson’s r = 0.60, 0.68, and 0.77 for stations 2, 8, and 5), which also indicated increased alpha diversity in denser, colder, and deeper water masses (Fig. 2B; Sup. Fig. 1; Sup. File 1, GoM AlphaDepth tab), and implied increased genome size is associated with increased G+C% content and alpha diversity, which corroborated previous studies.

Based on taxonomic annotations of the metagenomic reads against publicly available genomes (Kraken2 and Bracken results), the largest proportion of the microbial community was taxonomically classified from the surface and ML samples taken from stations 5 and 8 (∼20% vs <10%) and no more than 25% of the microbial community was classified in any sample (Fig. 2F), revealing that the GoM water column communities are under-represented in cultured-isolate databases. While samples EN21, EN84, and EN56 appeared the least similar to the other samples in their depth layers in terms of taxonomic composition (Fig. 2F-H), short-read, kmer-based beta-diversity clustering showed more similarity by depth layer than by sampling station (Fig. 2G-H). This depth-layer similarity held true even when GoM samples were compared to publicly available metagenome samples from other ocean basins (Sup. Fig. 2; Sup. File 1, Ocean Overview tab). Interestingly, alpha diversity was significantly lower in the GoM compared to the North and South Atlantic and Pacific samples (p < 0.05; Sup. File 1, Ocean Basin tab).

Members of the *Prochlorococcus* and *Pelagibacter* genera had the greatest abundance of any genera across all samples (12.5% and 5.3% max relative abundance, respectively) with greatest abundance in samples from the DCM to the surface (> 1% relative abundance DCM and above vs. < 1% below; Fig. 2F; Sup. File 2, Genus tab). *Prochlorococcus* was mostly absent below the DCM (< 0.1%) while some *Pelagibacter* species remained prominent members of deeper water communities although their relative abundance was much lower below the DCM (0.1-0.7%; Fig. 2F; Sup. File 2). The relative abundance of *Prochlorococcus* and *Pelagibacter* species were greatly reduced in sample EN21 compared to other samples above the DCM (<0.2%). In fact, the taxonomic abundance profiles for the surface and ML samples at station 2 looked quite different compared to stations 5 and 8 but the DCM, aOMZ, and OMZ profiles looked highly similar across all stations (Fig. 2F; Sup. File 2) indicating a signal from coastal proximity and/or that the freshwater riverine output from the Mississippi has a large influence on microbial assemblages of the ML and surface at this station.

Members of the *Pseudoalteromonas* and *Alteromonas* genera were the next most abundant across all samples with samples EN51 and EN84 showing the largest proportions of *Pseudoalteromonas* members (3.9% and 2.2%) and samples EN59, EN57, EN52 showing the greatest proportion of *Alteromonas* (1.5%, 0.8%, and 1.1%; Fig. 2F; Sup. File 2). Apart from sample EN84, which showed a high relative abundance of *Pseudoalteromonas* (2.2%), *Pseudoalteromonas* and *Alteromonas* species were greatly reduced in samples from station 2 and 8 (< 0.1%). They were also reduced in the DCM, aOMZ, and OMZ communities. *Pseudoalteromonas* abundance was greatest at the surface of station 5.

The taxonomic profile of sample EN56 was distinct from the other deep-water samples and all samples in general. Sample EN56 harbored a large proportion of members from the *Fonsibacter* (4.1%), *Nanopelagicus* (1.1%), and *Polynucleobacter* (0.8%) genera whose relative abundance was much lower in other samples (<0.01%) (Sup. Fig. 3 and 4). Consistent with these Kraken-based results, we recovered several rMAGs from sample EN56 matching these species such as *Candidatus* Fonsibacter ubiquis and *Candidatus* Nanopelagicus limnes (Sup. File 4). Sample EN56 was also the only sample below the ML with a high relative abundance of *Synechococcus* (0.3%) which was predominant in all surface and ML samples (0.3%-0.9%) but greatly reduced in DCM, aOMZ, OMZ and deep samples (<0.05%). Sample EN56 also had the largest fraction of classified reads of any sample below the ML (13.7% classified vs 7.8-10.3% in the DCM, 4.6-7.6% aOMZ, ∼5% OMZ and deep; Fig. 2F; Sup. File 2).

*Vibrio* abundance was greatest at the surface of station 5 (2.0%), but also the ML, aOMZ and deep samples EN52, 54, 58 and 59 (0.1-0.3%) and members of this genus were present but reduced in the remaining samples (0.03-0.07%; Fig. 2F; Sup. File 2). Members of the *Paenibacillus* genus were consistently abundant community members across all samples (0.13-0.25%). Similarly, members of the *Nitrosopelagicus* (*Thaumarchaeota*) genus were also consistently present at low abundance across all samples (0.001-0.003%) except in sample EN56. Members of the *Nitrosopumulus* (*Thaumarchaeota*) genus became abundant at and below the DCM (0.001-0.005% at the surface to 0.12-0.43% DCM and below). Relative abundance of *Acinetobacter* was highest at station 8 surface to aOMZ and station 2 ML (0.09-0.28%) but maintained a presence in all samples (0.04-0.08%) with lowest abundance in sample EN56. *Thioglobus* species were also consistently present in microbial communities across all samples (0.01-0.26%) with greatest abundance in the aOMZ and OMZ of all three stations, lower abundance in the surface and ML, and lowest abundance at the surface of station 2 (sample EN21) near the coast and Mississippi river outlet. Interestingly, the relative abundance of the bacteriophage *Eurybiavirus* was significant higher in the surface to DCM samples (0.02-0.50% vs. < 0.01% below the DCM, p < 0.05) with the greatest relative abundance in the ML at station 5. See figure 2 and supplemental file 2 for more extensive taxonomic details.

### Functional Gene Content Analysis

To assess sequence conservation and diversity at each station (vertically) or within each depth layer (horizontally) captured in our GoM metagenome samples we clustered CDS predicted from assembled contigs at 90% and 95% nucleotide sequence identity and 40% and 70% amino acid sequence identity (Fig. 3A-C) for various sample groupings at each depth layer (surface, ML, DCM, aOMZ, OMZ, and deep), each station (2, 5, 8) and all stations (surface-OMZ, and all depths). We chose 40, 70, 90 and 95% cutoffs because they are commonly used thresholds that roughly correspond to distinct taxonomic ranks (species, genus, and below genus) and provide a more and less conservative perspective of the data. If a gene cluster contained a CDS sequence from all samples in a grouping it was counted as a shared gene, if it contained CDS sequences from two or more (but not all) samples in a grouping it was counted as a flexible gene, and if it was a singleton (cluster of 1 CDS sequence only) or contained CDS sequences from only a single sample it was counted as a station-specific gene. This analysis showed that up to 35% of genes (OMZ layer 90% nuc and 40% and 70% aa clusters) were shared horizontally across all three samples taken from the same depth layer across the different stations (mean ∼24%) but only up to 4% of gene sequences were shared vertically across the top 5 depth layers taken from the same station (mean ∼3%; Fig. 3A; Sup. File 3). The surface and deep layers showed the fewest horizontally shared gene sequences likely due to samples EN21 and EN56 appearing as outliers in other analyses. The aOMZ layer, which contained sample EN84, was also an outlier and showed slightly fewer shared genes within this depth layer than the ML, DCM or OMZ layers (Fig. 3A; Sup. File 3). Interestingly, this analysis highlighted that 67% of genes across all samples contained nucleotide CDS sequence that is station-specific at the 95% sequence identity level (Fig. 3C; Sup. File 3), revealing the vast sequence diversity the oceans harbor and that our sequencing efforts are far from discovering all of it. Indeed, even clustering CDS sequences at 40% amino acid sequence identity, 60% of genes were found to be unique to a specific sample (i.e., station-specific), and the greatest percentage of shared genes between any two samples was 56% (shared + flexible genes) from the 90% nucleotide sequence identity gene clusters (Fig. 3A-B; Sup. File 3).

**Figure 3.**
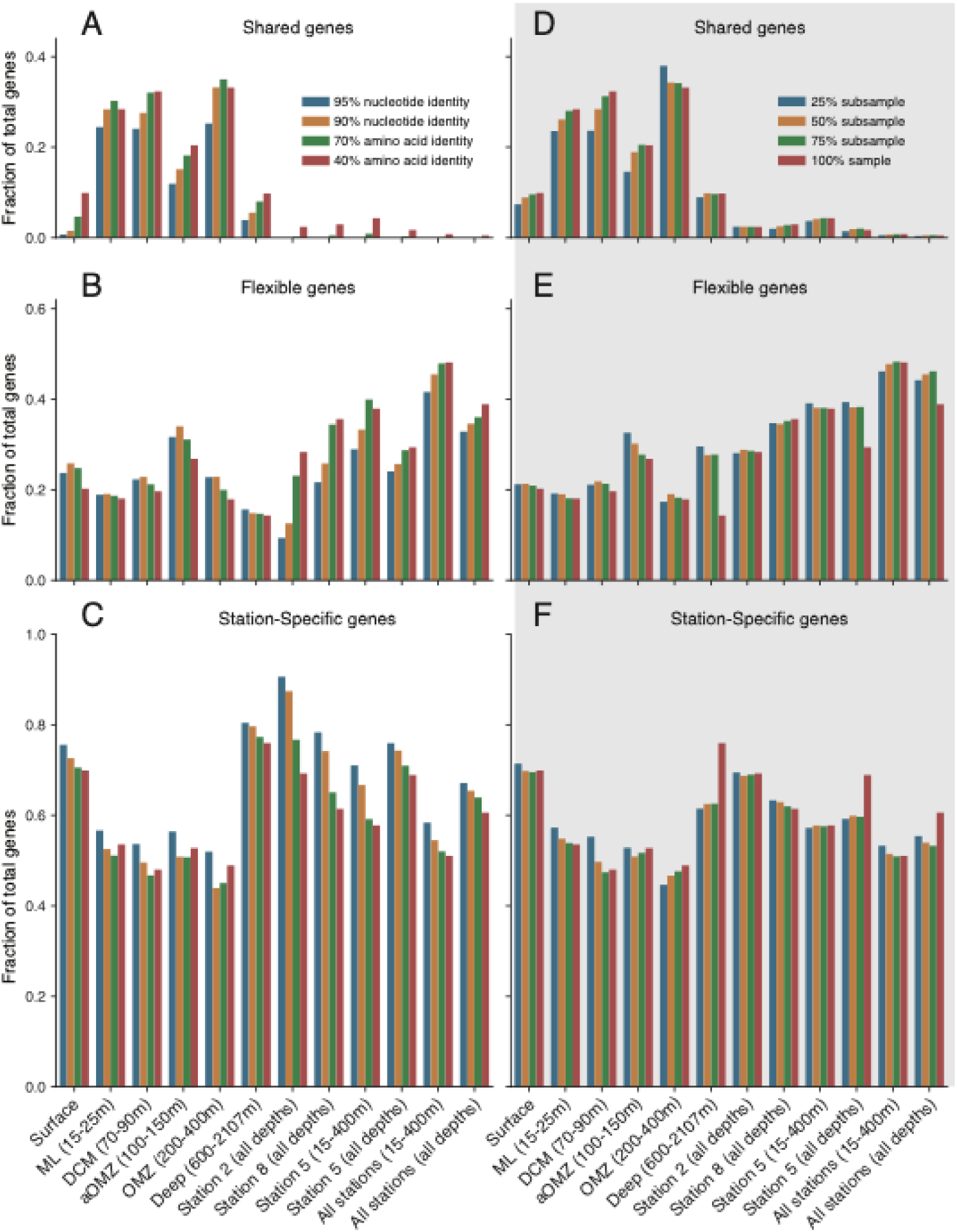
Gene sequence similarity by depth and by sample. Gene clustering results are partitioned into (A & D) genes shared by all samples in category (shared genes), (B & E) genes shared by two or more samples but not all (flexible genes), and (C & F) genes found in only a single sample (station-specific genes) in that category. Panels A-C display gene clustering results at 95% nucleotide sequence identity (blue bars), 90% nucleotide sequence identity (gold bars), 70% amino acid sequence identity (green bars), and 40% amino acid sequence identity (red bars). Panels D-F display gene clustering results from 40% amino acid sequence identity but for genes predicted from assemblies of down-sampled metagenomes at 25% subsample (blue blars), 50% subsample (gold bars), 75% subsample (green bars), and the full sample (red bars – same as A-C red bars). The various sample groupings are listed at the bottom along the x-axis with horizontal groupings by depth in the first six positions followed by vertical groupings by sampling station and all stations. Note the larger y-axis values in C and F indicating that the majority of assembled genes were station-specific. CDS were predicted with Prodigal from all assembled contigs for each sample and clustered with MMSeqs2.

To assess how much the sequencing effort or lack of complete coverage of sequencing diversity influenced the shared proportion of genes between samples, we subsampled each metagenome by randomly selecting without replacement 25%, 50% and 75% of the total reads. We then assembled the subsampled metagenomes, predicted CDS, and clustered the genes at 40% amino acid identity within each subsampled set separately (Fig. 3D-F; Sup. File 3, Subsampling tab). This analysis showed that the greatest difference, which was found between the 25% subsampled and full set (100% sample), was only an 8.71% reduction in shared genes. This result suggested that the ratio of shared to specific genes scales with sequencing effort and that we should expect similar results to those reported above even if we sequenced more deeply (but perhaps not as deep as to covering >99% of the estimated sequence diversity of the samples, which nonetheless was estimated to require an average of 92 ± 31Gbp to achieve based on Nonpareil analysis). This also aligned with our observations from sample EN56 which showed that station-specific gene sequence recovery also scales with sequencing effort.

Functional analysis based on COG categories and classes showed that the functional distribution of genes at broad category levels was similar (∼1% variance) across all stations and depths except for the Metabolism 1 class and the E category (Amino acid transport and metabolism) which had variance of 9.21% and 2% (Fig. 4; Sup. File 3, COG tab). The Mobile (X category) class genes had the least variance but also made up the smallest fraction of genes (2% or less). Samples EN21 and EN56 had the greatest proportion of hypothetical (“n/a” category), conserved hypothetical (S category – function unknown), and Mobile (X category) class genes and the smallest proportion of Metabolism 1(C, G, E, F, H, and I categories) and Metabolism 2 (Q and P categories) class genes out of all samples (Fig.4; Sup. File 3, COG tab). Of the Metabolism genes, samples EN21 and EN56 had the smallest proportions of categories P (Inorganic ion transport and metabolism), I (Lipid transport and metabolism), E (Amino acid transport and metabolism), and C (Energy production and conversion) genes out of all samples. Lastly, there were approximately 50,000 genes per sample on average (∼22% of predicted CDS) that did not receive an annotation and were not represented in the COG analysis. Sample EN21 had the greatest proportion of unrepresented genes at 38% which is 8% more than the second ranking samples in un-annotated gene counts (Sup. File 3, COG tab).

**Figure 4.**
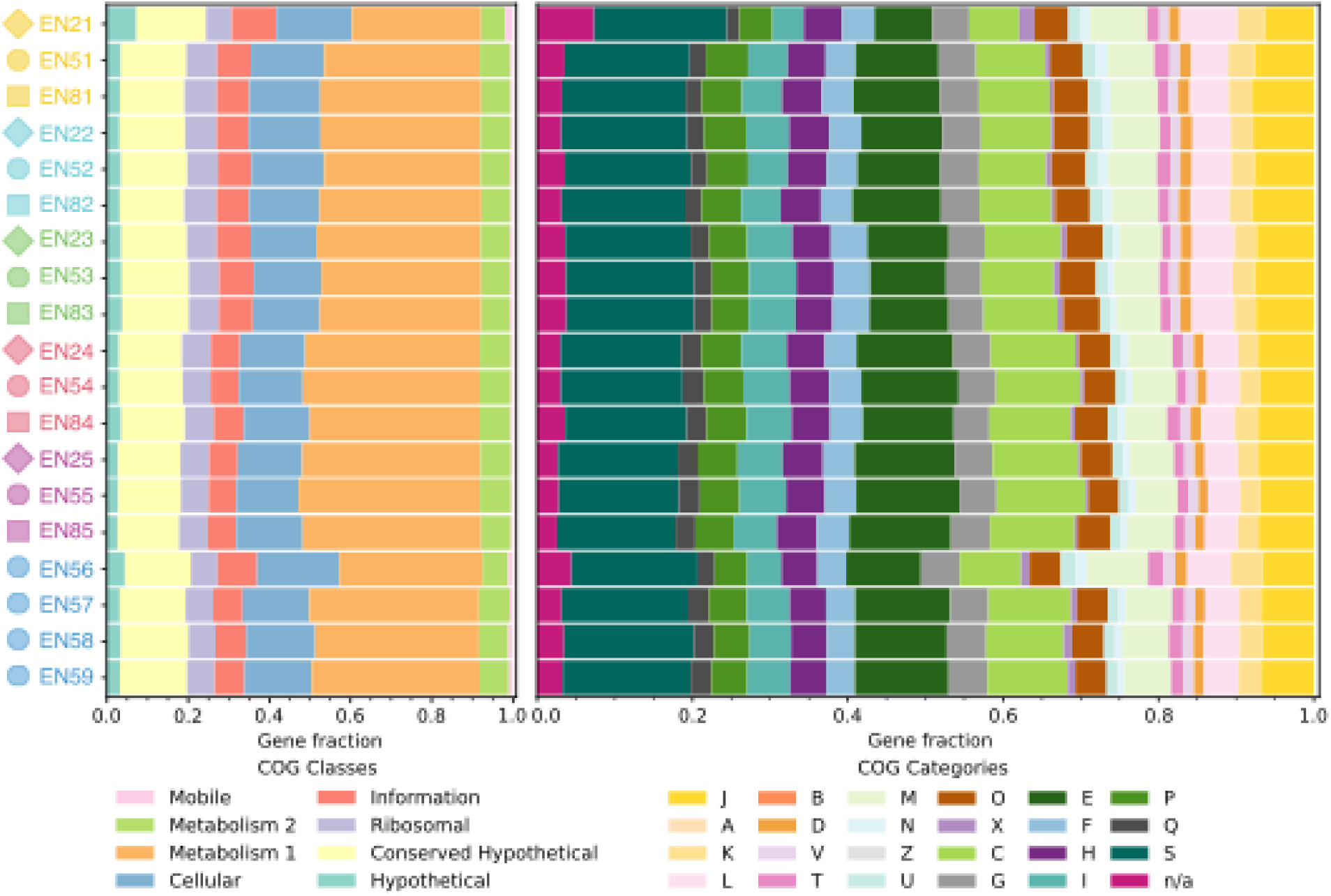
Distribution of COG major functional categories in each sample. Predicted CDS from assembled contigs of each metagenome were annotated with EGGNOG mapper. COG Categories (left) and COG Classes (right) were summarized to assess high level functional information. Note: gene fraction includes annotated genes only. 17-38% (mean 22%) of genes predicted for each sample did receive an annotation from EggNog mapper (i.e. did not find a good match in the database).

To probe deeper, we focused on gene sequences that were shared between all samples from the surface to the deep at station 5 and that received a non-hypothetical functional annotation. We mapped reads from each metagenome to each individual gene sequence originating from that metagenome and calculated a normalized relative abundance (see methods) for each gene in each sample (Fig. 5, “Single Genes” as blue circles). Next, we summed relative abundance of all genes assigned the same gene function for each sample (Fig. 5 “Summed Genes” as red circles). We computed correlations of summed genes against density, which was strongly correlated with temperature and depth (Sup. File 3, Gene Counts, Abundance and Correlation tabs). The 7,508 shared gene sequences from station 5 were assigned to 2,544 gene functions. Of these, 1,108 functions showed strong positive correlation (Pearson’s r ≥ 0.5) with density. An addition 339 functions showed strong negative correlation (Pearson’s r ≤ 0.5) with density, and the remaining 1,097 functions showed weak or no correlation (−0.5 < Pearson’s r < 0.5). This analysis showed 86 distinct dehydrogenase genes and 27 oxidoreductase genes that were positively correlated with depth (Pearson’s r ≥ 0.5; Fig. 5C,I). Likewise, it showed 21 distinct dehydrogenase genes and 3 oxidoreductase genes that were negatively correlated with depth (Pearson’s r ≤ −0.5; Fig. 5F). Additionally, there were 62 positively and 9 negatively correlated CoA related genes, different 50s ribosomal proteins (Fig. 5A,D), and phosphofructokinases (Fig. 5B,E), as well as beta-lactamases and transposases that showed increased abundance in colder, deeper water masses (Fig. 5G-H). Qualitatively, increased relative abundance of a function commonly coincided with increased genes of the function in the community (i.e. more genomes or species had distinct copies of the same function) but there were also examples of one or only a few gene sequences (or alleles) greatly increasing in abundance (i.e. particular community members carrying this gene became more abundant). We did not report further here on differentially present/absent gene sequences between the surface and the deep because relative abundance of these genes is below detection in some samples and includes over 430,000 gene sequences from station 5 alone, where 304,072 gene sequences (69% of all CDS from station 5) were specific to a single sample (Fig. 3; Sup. File 3, Gene Similarity tab), many of which were of unknown/hypothetical function (Fig. 4; Sup. File 3, COG tab).

**Figure 5.**
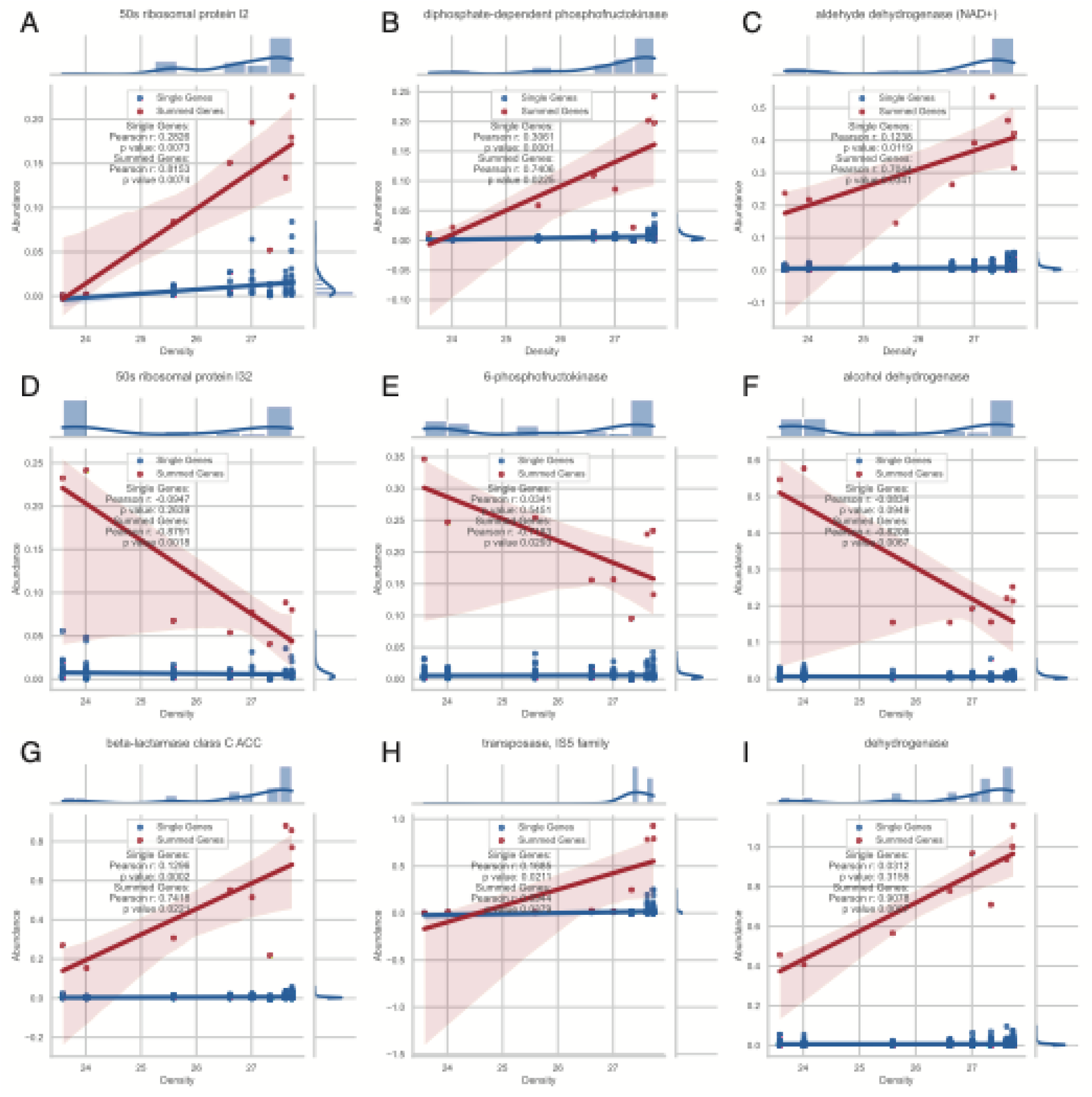
Selected gene functions that correlated with density at station 5. Normalized relative abundance (y-axis) is plotted against the sample’s density (σt, x-axis) for select gene functions (A-I) showing values of all individual genes assigned the stated function (blue circles) and the sum of those genes (red circles) for each sample. The corresponding best fit linear trend line with 95% confidence interval (shaded region) and correlation for each category (single genes or summed genes) is also plotted. Note that there were 9 samples collected at station 5 from the surface (density = 23.58, temperature = 27.18 C, depth = 3 m, far left points on x-axis) to the deep (density = 27.73, temperature = 4.25 C, depth = 2107 m, far right points on x-axis) and that depth and temperature correlated with density. For each gene, normalized relative abundance was calculated as TAD80/GEQ where TAD80 is the 80% truncated average sequencing depth (20% trimmed mean) and GEQ is genome equivalents as reported by MicrobeCensus. Functional annotations for each gene sequence were propagated from the representative sequence of the 40% amino acid gene cluster it was assigned.

### Description of rMAGs

In total we recovered 209 rMAGs from our GoM metagenome samples of which 154 were good quality with completeness ≥ 50% and contamination ≤ 10%. Of these, 145 shared < 95% AAI compared to MiGA’s NCBI Prok and TARA MAGs databases indicating that they likely represent novel species. About half of these MAGs have a match of >95% ANI against the GTDB reference genomes (n=72), revealing that related genomes have been recovered by previous studies but remain not-yet-named (Sup. File 4, rMAGs tabs). Interestingly, 168 of 209 rMAGs (80%) were assembled from only a single sample (Sup. File 4, Genomospecies tab) and 67 of 209 rMAGs (32%) were detectable (relative abundance > 0) in only a single sample (Sup. File 4, Detection tab). Further, 59 out of the 67 rMAGs detected in a single sample were detected in sample EN56 and 8 were detected in sample EN21. The remaining rMAGs exhibited 7 distinct patterns in relative abundance across the sample set (Fig. 6). Some rMAGs were detectable in the surface and ML or only the DCM while others were detected in the surface, ML, and DCM. Similarly, some rMAGs were detected in only the aOMZ and OMZ while some were detected in the aOMZ, OMZ, and deep or the DCM, aOMZ and OMZ. Only 4 rMAGs were detected in the deep only. No rMAGs were detectable in all depth layers or all samples, but a couple were detectable in both deep and shallow waters, revealing remarkable versatility across the water column. The latter MAGs included *Alteromonas macleodii* (97% AAI to the most closely related genome available), and *Desulfuromonas sp.* (40% AAI to *Desulfuromonas soudanensis*), which showed relatively high abundance at both the surface (0-200m) and the deep (>1000m) (Fig. 6).

**Figure 6.**
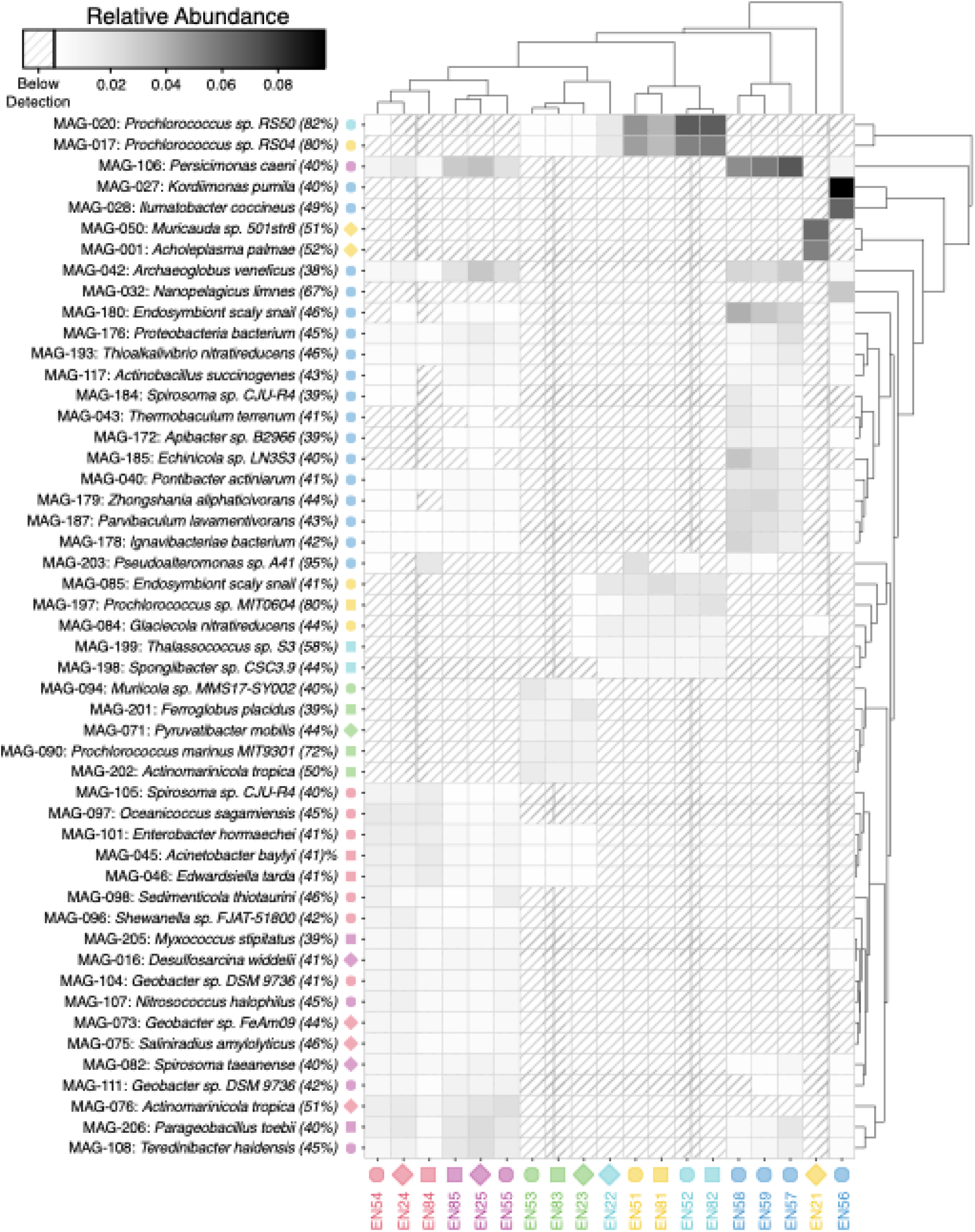
rMAG Relative Abundance in each sample. Relative abundance calculated as TAD80/GEQ. Genome equivalents (GEQ) computed with MicrobeCensus. Top 50 rMAGs with greatest cumulative abundance are reported on the rows. The MAG number links to additional information in Supplemental Excel File 3. Taxonomy assignment is closest matching genome in MIGA’s NCBI Prok database and the percentage is the AAI of the rMAG to the closest match. Color and shape indicate sample of rMAG origin. Columns are metagenome samples. Note that apart from sample EN21, samples cluster by depth. Hierarchical clustering computed with the scipy.cluster.hierarchy.linkage() function from the SciPy package in Python with method=’average’ to calculate cluster distance using the UPGMA algorithm.

## DISCUSSION

Collectively, our results show that a large proportion of microbial diversity in the GoM is unknown to our genome and gene databases, and, even from the known genera or families detected by our metagenomics efforts, the rMAGs recovered within them appear to represent predominantly novel species. Similarly, only a maximum of 25% of the microbial community was taxonomically classifiable to represent a known (named) species at the read level (Fig. 2F), further corroborating that the GoM water column communities are under-represented in cultured- and culture-independent databases. Therefore, sequencing efforts to characterize even environments that are thought to be well-characterized by now such as the oceans, remain worthwhile towards cataloguing the extant diversity on the planet. It might be the case that the GoM harbors disproportionately more genome diversity than other oceans due to its proximity to major human populations and activities, but also uncommon effects of oil seeps and major rivers such as the Mississippi River. The fact that the GoM metagenomes show different alpha diversity than other previously characterized ocean basins such as the North and South Atlantic and Pacific oceans (p < 0.05; Sup. File 1, Ocean Basin tab), is also consistent with this interpretation.

Another highlight of our analysis is the somewhat surprising number of gene sequences specific to only a single sample. While it is expected to find more shared genes across similar depths (horizontally) than between depths (vertically), the amount of vertically shared genes is rather low and the proportion of station-specific gene sequences correspondingly high. For example, our analysis showed that 67% of genes across all samples represent unique sequences at the 95% nucleotide identity level (Fig. 3C; Sup. File3), and that the greatest percentage of shared genes between any two samples was only about 56% at this identity level (Fig. 3A-B; Sup. File3). Our projection is that even with 5- or 10-times greater sequencing effort, the amount of station-specific gene diversity would decrease by less than 10%. Consistent with these results, the 340% increase in sequencing effort for sample EN56 yielded relative to other samples did not result in proportions of sequence recovery or shared gene sequence outside the range of other samples, which also suggested that station-specific gene sequences scale with sequencing effort presumably due to increased recovery of sequence from the rare biosphere (Sup. File 1). These results somewhat echo the large genome-specific genes (e.g. >50% of total genes detected) revealed by the comparative analysis of genomes of isolates of the same species [e.g. (34)], revealing that the concept of the pangenome may apply equally well to whole communities, not only to the diversity within a species. Perhaps these results are attributable, at least in part, to the planktonic nature of marine microbes since the parameters of populations mixing and migration are controlled by currents where similar species occupying the same niche space drift apart in separate water parcels and diverge by neutral evolutionary processes. It’s also possible that the influence of these currents, water masses, seeps, and river inputs might be more exaggerated by the unique characteristics of the GoM relative to other ocean basins.

It is also notable that the effect of the Mississippi River (or others) on the GoM could be so pronounced that sample EN56 270 miles from the coast and 600m deep had a peculiar freshwater signal based on the relative abundance of microbial community members. This sample had a taxonomic profile distinct from the other deep-water (and all) samples with the top three most abundant species detected being *Candidatus Fonsibacter ubiquis* (4.1%), *Candidatus Nanopelagicus limnes* (0.8%), and *Polynucleobacter acidiphobus* (0.7%), all known to inhabit various freshwater or low-brackish conditions and not previously identified at higher salinity [(35–37); Sup. File 2, Species tab]. The relative abundance of these species in other samples was much lower (0.02 ± 0.01% *Fonsibacter*, and < 0.001% others). The 4^th^ most prominent member of the EN56 community was *Paenibacillus larvae*, a honeybee parasite (38), but relative abundance of this species was consistent with other samples (0.16 ± 0.03%). Sample EN56 was also the only sample below the ML with a high relative abundance of the coastal *Synechococcus* (0.3%), which was predominant in surface and ML samples (0.3-0.9%) but reduced in DCM, aOMZ, OMZ and deep samples (<0.05%), consistent with the hypothesis of a deep freshwater influence. Finally, the salinity measured for sample EN56 was in line with the other deep-water samples (∼34.9 PSU; Sup. File 1) which is far from the low-brackish and freshwater conditions these species have been identified in previously. Of course, with peculiarities such as this, bottle leakage or some process of surface water mixture or sinking could also occur, but we did not note any evidence of bottle linkage and the community signal from station EN56 was quite distinct from our surface samples. Thus, this signal is highly unlikely to be attributed to contamination and it could be an interesting line of investigation in future research.

Another sample, EN21 approximately 50 miles from the Mississippi River outlet, exhibited a freshwater signal as well, although its signal was more evident in the physicochemical measurements than the identification of freshwater associated species. The surface at station 2 showed lower salinity (29.6 vs. 36 PSU) and increased fluorescence (0.9 vs. 0.07 mg/m^3^) with a reduced DCM layer fluorescence peak (0.3 vs. 0.8 mg/m^3^), and the taxonomic profiles of the surface and ML were different, containing a greater proportion of unclassified reads and reduced relative abundance of *Prochlorococcus* (Figs. 1B-D & 2F) compared to stations 5 and 8. The surface sample at station 2 also had reduced relative abundance of *Pelagibacter* species. This finding reveals a brackish water community in the surface ocean surrounding the Mississippi River (and likely others) that is largely unknown in current databases and needs to be explored further in future research. However, since the prominent freshwater species detected in sample EN56 are not reflected in sample EN21 or 22, this brackish water community identified at station 2 does not explain why we detected freshwater species so far offshore in salt water, and below the OMZ in sample EN56 at station 5.

We also find it interesting that the MAGs from samples below the DCM seemed to bin easier, or at least represent a greater proportion of the community, even though the deep metagenomes harbored larger genomes and gene/sequence diversity. One possible explanation for this is that the surface harbors species that have high intra-species sequence diversity, or such species make a larger fraction of the total community, such as *Prochlorococcus* and *Pelagibacter*. Our recent work based on Single-cell Amplified Genomes (SAGs) has confirmed the high intra-species diversity for deep-sea SAR11 genomospecies, e.g., intra-species ANI ranging between 91 and 100% vs. 96-100% for most, well-sequenced species (33). High intra-species diversity is not handled well by most, if not all, pieces of software for assembly and/or binning e.g., assemblers are tuned to merge sequences that are 97-98% identical at the nucleotide identity or higher (39).

Several of our results also corroborated previous findings, but with interesting additions and/or expansions. For example, we observed an increase in genome size and the abundance of certain genes with depth. However, we also detected a distinct signal in a sample located between the OMZ and the deep layer (sample EN56, mentioned above). Our data also suggests that genome size and other parameters correlate more strongly with density than with depth or temperature alone, indicating that water mass boundaries may play a greater role than pressure or temperature. Additionally, some gene functions that increase with depth are paralleled by similar but distinct functions that decrease with depth—for instance, aldehyde dehydrogenase versus alcohol dehydrogenase, or diphosphate-dependent phosphofructokinase versus 6-phosphofructokinase (Fig. 5C vs. F and B vs. E). Functional analysis based on COG categories and classes showed that the Metabolism 1 class and the E category (amino acid transport and metabolism) had substantial variance with depth. While similar reports of gene functions increasing in the deep have been reported previously (23–25), we find it interesting that many of these gene functions have similar alternate gene functions that are more abundant at the surface which could indicate possible sequence adaptation to the unique physicochemical properties of the surface vs. deep waters and/or functional differentiation (Fig. 5). Another interesting observation is that increased abundance of a gene frequently corresponded to more distinct copies of those genes found in the community (i.e., being community-wide) versus an increase in abundance of one or few species carrying that gene (or a specific allele, defined at the >95% nucleotide identity level). A few previous studies suggest that these gene differences could represent important physiological adaptations associated with ocean depths changes in temperature and pressure, such as differential presence of ribosomal proteins involved in catalyzing peptide bond formation (40, 41), or differences in lactate dehydrogenase function (42).

To conclude, there remains much to be learned about the ocean microbiome, and bioinformatics sequence analysis alone is not enough. The overwhelming number of unknown taxa and gene functions require specific and dedicated efforts to uncover these functions, yet there is still much to be gained from continued sequencing efforts. Understanding genetic differences between species, and between divergent populations of the same species, requires data on population tracking, migration, and recombination. Sampling efforts should target distinct water masses and the boundaries between them as well as incorporate ocean current models into sampling schemes. Our work here provides a detailed quantitative analysis of microbial communities in the GoM water column sampled while tracking *Trichodesmium* blooms tracked by satellite imaging on the surface during a period without major oil inputs. It also uncovered some interesting signals to explore and compare against in future research endeavors.

## METHODS

### Sample collection, DNA extraction and sequencing

Samples were collected from three depth profiles of the Gulf of Mexico (GoM) on 29 May 2012 aboard the R/V Endeavour (cruise EN509; https://www.bco-dmo.org/dataset/4067) at station 2, 5 and 8, representing surface (51, 21, 81), mixed layer (52, 22, 82), deep chlorophyll maxima (53,23, 83), above oxygen minimum zone (54, 24,84), oxygen minimum zones (55, 25, 85), and bathypelagic water masses (Sup. File 1). Collections were made using Niskin bottles on a rosette containing a conductivity–temperature–depth profiler (Sea-Bird SBE 911plus).

Water samples (20 gallons per sample) were pre-filtered through a nylon disk filter (47mm, 1.6 um porosity, Whatman, GE Healthcare Bio-sciences, Pittsburgh, PA, USA), and biomass was collected on a glass fiber disc filter (47 mm, 0.2 μm pore size, Vendor) via a peristaltic pump. Disc filters were immediately frozen after collection at −80oC and DNA extractions were performed back in the laboratory.

DNA was extracted from the disc filters using an enzymatic lysis and phenol protocol as previously described (Tsementzi et al., 2014). Briefly cells were lysed with the addition of lysozyme (1 mg/μl final concentration in 5ml of lysis buffer per filter) and incubation for 30 min at 37oC. Addition of Proteinase K (1 mg/100 μl lysis buffer with 100 μl 20% SDS) was followed by a 2h incubation at 55oC. DNA was extracted by phenol:chloroform extraction, followed by ethanol precipitation and wash. DNA was purified using the Ampure XP-Beads (Beckman Coulter).

### Metagenome QC, Assembly, and Binning

Paired Illumina reads for each metagenome sample were trimmed and quality filtered with BBDuk with parameters “qtrim=w,3 trimq=17 minlength=70 tbo=true tossjunk=t cardinalityout=t.” The trimmed and filtered read set for each sample (TRIM) was also normalized (NORM) with BBNorm with parameters “target=30 min=5 prefilter=t.” BBDuk and BBNorm versions were part of the BBMap v38.93 release from 09/21/2021 (43). For each sample, both the TRIM and NORM read sets were assembled separately with IDBA-UD v1.1.3 with default parameters (44). Assembled contigs <1000 base pairs in length were removed. For each sample, both the TRIM and NORM assemblies were binned separately with MaxBin v2.2.7 with default parameters and MetaBAT v2.12.1 with default parameters (45, 46). This generated 4 overlapping metagenome assembled genome (MAG) sets for each metagenome sample (maxbin_trim, maxbin_norm, metabat_trim, metabat_norm), which were dereplicated using the “derep” workflow from MiGA v1.0.0 to select the highest quality MAG from each 95% ANI cluster within each sample (47). The dereplicated MAGs from each sample were then further dereplicated across all samples creating the GoM representative MAG set (rMAGs). The quality and taxonomic classification of each rMAG was assessed using the “quality” and “classify” workflows from MiGA as well as the “lineage” workflow from CheckM v1.1.3 with default settings and the “classify” workflow from GTDB v1.7.0 with default settings (48, 49). Metagenome data provided in supplemental file 1 and rMAG data provided in supplemental file 4. The TRIM read sets and contig assemblies were used for all other downstream analyses.

### Predicted CDS and Functional Annotation

Genes were predicted for all assembled contigs using Prodigal v2.6.3 with setting “-m meta” for metagenomes (50). Functional gene annotations for the relative in situ abundance and density correlation analysis was performed with MicrobeAnnotator v1.0.4 with default settings, Diamond v2.0.1 and Kofam, Swissprot, Trembl, and RefSeq databases created February 10^th^ 2021 (51, 52). Functional gene annotations for the high-level COG class and category analysis was performed with EggNog mapper v2.1.9 with default settings with EggNOG 5 database (53, 54).

### Metagenome Sample Diversity

Alpha diversity and sequence coverage were estimated with Nonpareil v3.401 with settings “-T alignment -f fastq -× 100000” using only the forward read pair from each sample (55). Beta diversity analysis was performed with Simka v1.5.3 with default settings (56). Community taxonomic distribution was estimated with Kraken v2.1.2 and Bracken v2.5 with default settings (57, 58). Gene similarity between metagenome samples was assessed by gene clustering at 95% and 90% nucleotide sequence identity and 70% and 40% amino acid sequence identity using MMSeqs2 v13.45111 with settings “--cov-mode 1 -c 0.5 --cluster-mode 2 --cluster-reassign,” and “--min-seq-id” of 0.95, 0.90, 0.70, or 0.40 respectively (59). A gene was considered shared if a CDS from all samples in the grouping was found within a gene cluster, flexible if it was found in two or more samples in a grouping, and station-specific if was found in only one sample in the grouping. Groupings are described in Fig. 3 and in the text.

### Relative Normalized Abundance Estimates

Average genome size and genome equivalents (GEQ) were estimated with MicrobeCensus v1.1.0 with setting “-n 100000000” to use all reads (60). A separate database was used for assembled contigs from each sample to map reads for the functional gene normalized abundance estimates, and a single database was used containing all rMAGs to competitively map reads for rMAG relative normalized abundance estimates. Databases were created using the makeblastdb command from Magic-Blast, and reads were mapped to the databases with Magic-BLAST v1.5.0 with settings “-infmt fasta -paired -no_unaligned -splice F -outfmt tabular -parse_deflines T” (61). Read mapping results were filtered to retain only the single best hit for each read and to remove alignments with less than 90% match length (calculated as alignment length / read length) and any reads less than 70 base pairs in length. Relative normalized abundance was calculated as TAD80/GEQ where TAD80 is the truncated average sequencing depth discarding 10% of base pair positions with the highest and lowest read mapping depth (retaining 80%) before computing the average depth (depth at each base pair position / length of positions) to remove any outlier positions that may skew the mean. TAD80, which is the sequencing depth, is normalized by GEQ.

### Additional Analysis and Figures

Apart from the supplemental statistics presented in the Excel files, all other analyses and figures were generated using in house Python scripts prepared for this study. Scripts were written for and executed with Python version 3.7.10+ and used the following packages: Matplotlib, Seaborn, Scipy, Statsmodels, Pandas, Numpy, and Seawater (62–69). All code has been deposited in the GitHub repository created for this manuscript.

## ACKNOWLEGMENTS

We want to thank PACE (The Partnership for an Advanced Computing Environment) at Georgia Tech for providing computational resources. This work was supported, in part, by US NSF Awards 1831582 and 2129823 to K.T.K. JM was supported by a grant from the Gulf of Mexico Research Initiative to support the ECOGIG (Ecosystem Impacts of Oil and Gas Inputs to the Gulf) Consortium.

## COMPETING INTERESTS

Authors declare they have no competing interests.

## CODE AND DATA AVAILABILITY

Metagenome sequences deposited to NCBI SRA associated with bioproject PRJNA291283. Accession number for each sample listed in supplemental file 1 – GoM Overview tab.

Code and rMAGs available at: https://github.com/rotheconrad/Descriptive_Metagenome_GoM Zenoda DOI: https://doi.org/10.5281/zenodo.15214590

